# DIISCO: A Bayesian framework for inferring dynamic intercellular interactions from time-series single-cell data

**DOI:** 10.1101/2023.11.14.566956

**Authors:** Cameron Park, Shouvik Mani, Nicolas Beltran-Velez, Katie Maurer, Satyen Gohil, Shuqiang Li, Teddy Huang, David A. Knowles, Catherine J. Wu, Elham Azizi

## Abstract

Characterizing cell-cell communication and tracking its variability over time is essential for understanding the coordination of biological processes mediating normal development, progression of disease, or responses to perturbations such as therapies. Existing tools lack the ability to capture time-dependent intercellular interactions, such as those influenced by therapy, and primarily rely on existing databases compiled from limited contexts. We present DIISCO, a Bayesian framework for characterizing the temporal dynamics of cellular interactions using single-cell RNA-sequencing data from multiple time points. Our method uses structured Gaussian process regression to unveil time-resolved interactions among diverse cell types according to their co-evolution and incorporates prior knowledge of receptor-ligand complexes. We show the interpretability of DIISCO in simulated data and new data collected from CAR-T cells co-cultured with lymphoma cells, demonstrating its potential to uncover dynamic cell-cell crosstalk.

**Availability:** DIISCO is publicly accessible at https://github.com/azizilab/DIISCO_public. All data will be deposited to GEO upon publication.

## 1 Introduction

Single-cell RNA sequencing (scRNA-seq) which measures gene expression at the resolution of individual cells is a powerful technology for elucidating heterogeneous cell types and states [25,28,2]. The recent expansion of single-cell datasets profiling biological systems and longitudinal clinical cohorts over multiple time points offers an exciting opportunity to characterize the temporal dynamics of cell types and cell-cell crosstalk. However, there is an unmet need for computational frameworks that can effectively integrate single-cell data across time points while accounting for temporal dependencies, especially in longitudinal clinical data where the timing of biopsies cannot be designed or controlled and is often extremely variable across patients [3].

Complex systems such as the tumor microenvironment are ever-changing, especially with disease progression or therapy. Various cancerous and non-cancerous cell types are engaged, and interactions between these cells lead to diverse treatment responses. Uncovering crosstalk between tumor and immune cells [14] for example can unravel immune dysfunction mechanisms. Furthermore, characterizing the dynamic nature of these interactions and their effect is crucial for understanding mechanisms underlying response or resistance to therapies and how this machinery can be leveraged to develop more effective therapeutic strategies [31,23]. Current approaches for predicting cell-cell interactions using scRNA-seq data rely on existing databases of interacting protein complexes, and they predict interactions based on expression levels of known receptor-ligand (R-L) pairs [10,6,11,16]. For instance, CellphoneDB infers interactions based on complementary minimum expression of protein subunits [10]. CellChat scores interactions based on differentially expressed genes and has been shown to reduce the number of false positive interactions [11]. NicheNet infers interactions through the expression of downstream target gene pathways for different R-L pairs [6]. Current methods, however, have several limitations. The receptor-ligand subunits can have variable effects and expressions in different cell types, a context-dependent nuance often overlooked by existing tools. Reliance on databases also limits the discovery of novel interactions and rare understudied cell types. Importantly, existing tools only predict static interactions. They are not capable of inferring dynamic time-varying interactions from time-series datasets and do not obtain the strength or sign of interactions (e.g. inhibitory or activating crosstalk).

Here, we present DIISCO (Dynamic Intercellular Interactions in Single Cell transcriptOmics), an open-source tool (https://github.com/azizilab/DIISCO_public) for joint inference of cell type dynamics and communication patterns. We define cell types or states as clusters of cells with similar gene expression profiles in scRNA-seq data. To model the temporal dynamics of cell types, i.e. changes in their frequency over time, while addressing the challenge of varying time intervals, we deploy Gaussian Process (GP) regression models [29]. GPs provide sufficient flexibility with nonparametric function learning, and therefore do not require any knowledge of form or rate of changes of cell types, thus expanding their applicability. They have been proven successful in integrating time-series single-cell datasets [18], and in identifying key cell states with distinct temporal dynamics shaping response to immunotherapy in human leukemia [3]. By utilizing GPs, which encode temporal dependencies between all pairs of time points, we expand the utility of the model to clinical settings and aggregating both short-term and long-term patient specimens.

We additionally draw inspiration from Gaussian Process Regression Networks (GPRNs) which have been previously applied to gene expression dynamics as well as datasets in finance, physics, geostatistics, and air pollution [29,17]. GPRNs are Bayesian multi-output regression networks that leverage the flexibility and interpretability of GPs as well as structural properties of neural networks. In this work, we use the GPRN concept to encode interactions between cell types. The ability to capture highly nonlinear and time-dependent correlations between outputs makes GPRNs ideal for learning complex cell-cell interactions and their variability over time.

DIISCO is a Bayesian framework that infers dynamic interactions in scRNA-seq data from non-uniformly sampled time points, incorporates prior knowledge on receptor-ligand complexes, and quantifies uncertainty in both its predictions and interpretations. Unlike previous methods, DIISCO can be trained on temporal data to learn how interactions change over time between samples. We demonstrate the performance of DIISCO on simulated data as well as new data collected from Chimeric antigen receptor (CAR) T cells interacting with lymphoma cells.

### Contributions

In brief, the key contributions of our work are: (1) Construction of a probabilistic framework for modeling temporal dynamics of diverse and interacting cell types in complex biological systems, through the integration of single-cell RNA-seq datasets collected across non-uniformly sampled time points; (2) Development of first-in-class computational tool for predicting time-resolved cell-cell interactions, along with the strength and sign (activating/inhibitory) of interactions, empowering discoveries of cell communication associated with progression of biological processes or treatments; (3) A novel Bayesian framework for incorporation of prior knowledge on signaling protein complexes with time-series single-cell RNA-seq data to improve identifiability of dynamic cell interactions and quantify uncertainty; (4) Demonstration of performance in simulated and new lymphoma-immune interaction data.

## 2 Materials and Methods

### 2.1 General Overview

DIISCO captures the temporal dynamics and interactions of cell types using scRNA-seq data from multiple time points. The time-series single-cell data are first merged to define unique cell types or states through typical clustering approaches [30,15]. The pooling of cells helps with improving statistical power in the detection of rare cell types. Then, the number of cells assigned to each cell type at each time point is computed and used as training data for DIISCO (**Fig. 1a**). The cell type dynamics can be in the form of cell counts at each time point or standardized proportions of each cell type (normalized by total cells in the sample) at each time point. Using proportions is often useful in complex patient data when the sampling and quality of biopsies are inconsistent over time points.

**Fig. 1.**
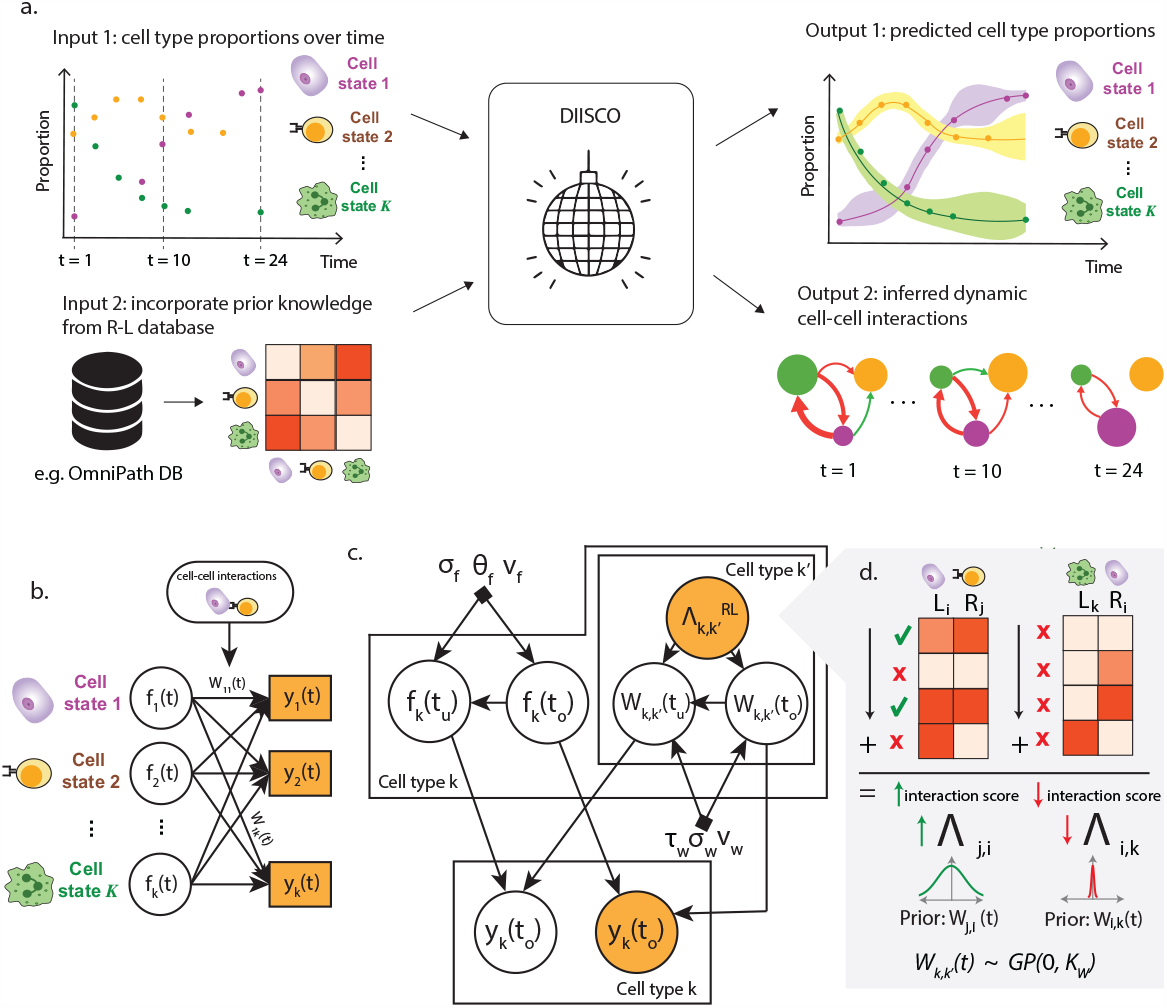
Overview of DIISCO framework. **a** General workflow of DIISCO algorithm including inputs and outputs. Cell type proportions are computed from scRNA-seq data in each timepoint. Expression of RL complexes is incorporated to obtain time-resolved interactions between cell types. **b** Network diagram of model framework. **c** Graphical model of DIISCO, including all hyperparameters. Latent variables are depicted as white circles, and observed variables are depicted as yellow-shaded circles. **d** Process for incorporating domain knowledge on receptor-ligand interactions as a prior for cell-cell interactions into DIISCO.

**Fig. 2.**
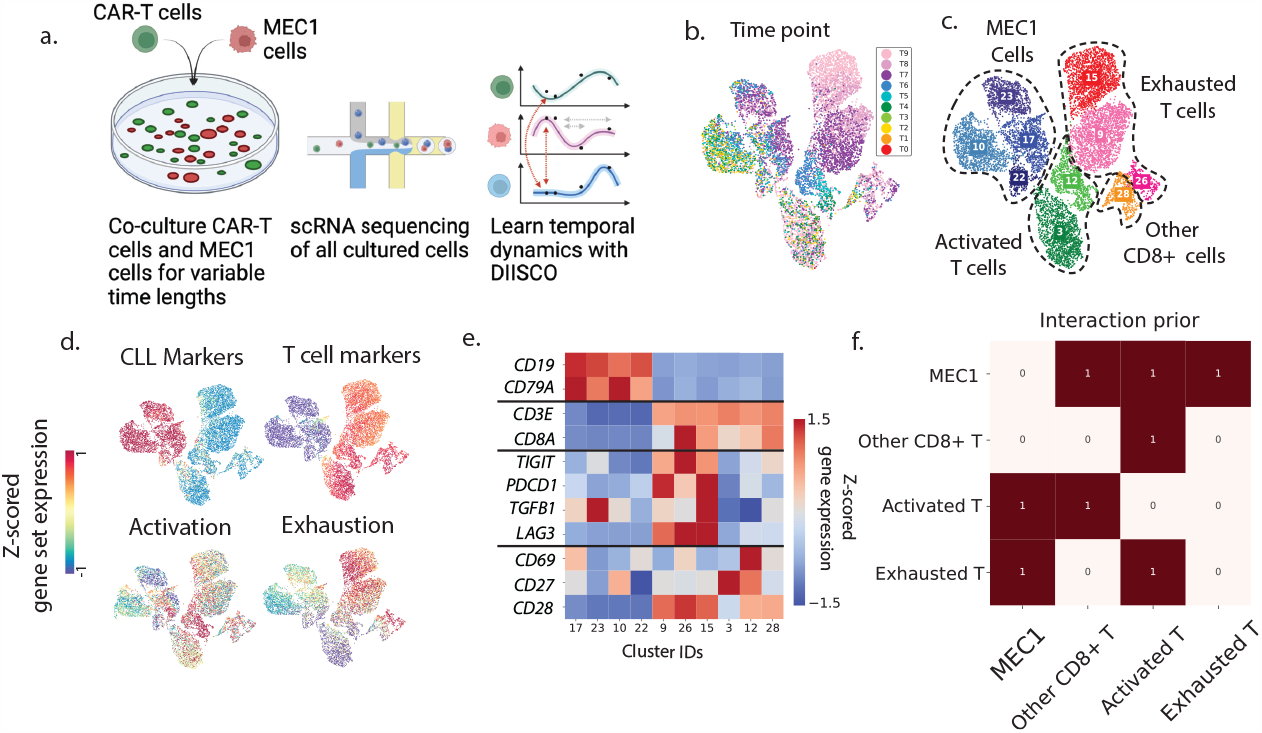
Experimental setup and data preprocessing. **a**. Overview of data collection process and analysis. **b-d**. 2D UMAP projection of 9082 single-cell transcriptomes from one co-culture experiment colored by time point (b), cluster and cell type assignment (c), and average expression of different gene set markers (d) used to define metaclusters. **e**. Heatmap of average cluster expression of individual genes used in gene set analysis, z-scored across all clusters. **f**. Prior constructed for DIISCO input, generated as explained in Section 2.3

DIISCO also utilizes an optional prior based on the expression of receptor ligands (RLs). Our framework learns interactions from both RLs and temporal dynamics of cell types, and reliance on RLs can be adjusted with the prior strength. Utilizing the RLs is beneficial for constraining the solution space of interactions and improving identifiability and model robustness. The output of DIISCO is two-fold. First, DIISCO outputs predicted cell type dynamics. This is helpful for assessing model fit, especially when tuning hyperparameters. Second, DIISCO outputs a time-series interaction matrix that reveals which cell-cell interactions are evolving with time (**Fig. 1a**).

### 2.2 Notations

We assume that we have *N* measurements of cell type frequencies at time points *t*_1_, …, *t*_*N*_. We define *y*(*t*_*i*_) as a *K*-dimensional vector of observations at time *t*_*i*_ where the *k*-th dimension corresponds to the frequency of the *k*-th cell type. Additionally, we assume we have a set of *M* unobserved time points *t*_*N*+1_, …, *t*_*N*+*M*_, placed anywhere on the time axis, for which we would like to infer the cell type values *y*(*t*_*N*+1_), …, *y*(*t*_*N*+*M*_). We denote the set of all time points as 𝒯= {*t*_1_, …, *t*_*N*+*M*_ }and call 𝒯_*u*_ the set of unobserved time points 𝒯_*u*_ = {*t*_*N*+1_, …, *t*_*N*+*M*_ }and 𝒯_*o*_ the set of observed time points 𝒯_*o*_ = *t*_1_, …, *t*_*N*_. Throughout the paper, we will use the convention of *·*_*u*_ and *·*_*o*_ to denote unobserved and observed variables respectively. In the remainder, for ease of exposition, we will refer to *y*(*t*_*i*_) as proportions with the understanding that either proportions or cell counts could be used, although the results should be interpreted in a different manner depending on the value used (see section 4).

Additionally, we have a binary matrix *Λ* of size *K×K* where *Λ*_*k,k*_*′* = 1 indicates that the *k*-th cell type might interact with the *k*^*′*^-th cell type and *Λ*_*k,k*_*′* = 0 indicates that they don’t. We allow for non-symmetric matrices to allow for directionality in interactions. In practice, we obtain this matrix from measurements of expression of known receptor-ligand complexes as discussed in the next section.

### 2.3 Model Specification

DIISCO is a generative model that assumes cell type frequencies, denoted as *ŷ* (*t*_*i*_), are derived from the following process (**Fig. 1b**):

For every time point, we sample a set of latent features, *f* (*t*_*i*_) *∈* ℝ^*K*^ where every feature, i.e every coordinate of *f* (*t*_*i*_), is a Gaussian process. We call this set *ℱ* = {*f* (*t*_*i*_) | *t*_*i*_ *∈𝒯*} and let ℱ_*o*_ and ℱ_*u*_ denote the set of latent features at observed and unobserved time points respectively.

Similarly, we sample at each time point an interaction matrix *W* (*t*_*i*_) *∈* ℝ^*K×K*^ where we also assume that each of the *K× K* coordinates of *W* (*t*_*i*_) is a Gaussian process across time. We call this set 𝒲 = {*W* (*t*_*i*_) | *t*_*i*_*∈𝒯*} and let 𝒲_*o*_ and 𝒲_*u*_ denote the set of interaction matrices at observed and unobserved time points respectively.

Finally, we sample the standardized cell type proportions *ŷ*(*t*_*i*_) *∈* ℝ^*K*^. We do this by sampling from a multivariate Gaussian with mean 𝒲 (*t*_*i*_)*f* (*t*_*i*_) and covariance 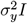. In other words:

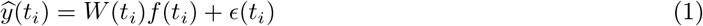

where 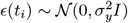 represents a zero-centered Gaussian noise process. We use *𝒴*= *{ŷ* (*t*_*i*_) | *t*_*i*_ *∈ 𝒯 }* to denote the set of all standardized cell type proportions across all time points with *𝒴*_*o*_ and *𝒴*_*u*_ having the same meaning as before ^13^.

Figure 1c depicts this process graphically ^14^ and Algorithm 1 details this process.

#### Algorithm 1

Generative Process used by DIISCO

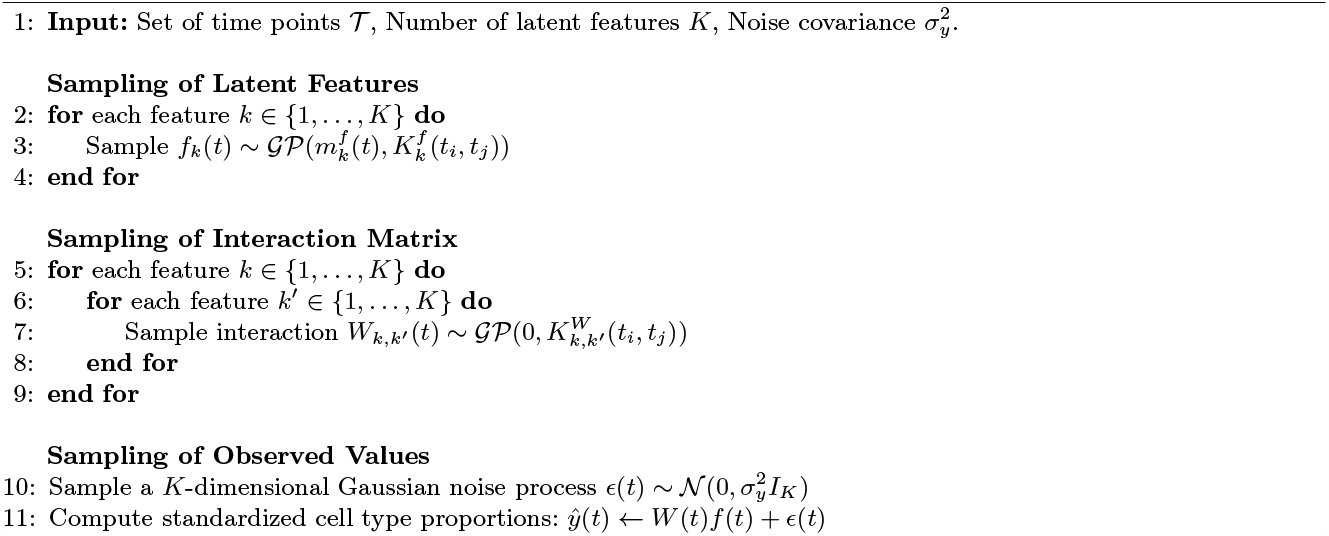

#### Model interpretation

The latent variable *W* (*t*)_*i,j*_ represents the direction and effect of intercellular interaction (signaling communication) from cell type *j* to cell type *i* at time *t*. In particular, *W* (*t*)_*i,j*_ *>* 0 represents an activating effect, *W* (*t*)_*i,j*_ *<* 0 denotes inhibitory impact, and *W* (*t*)_*i,j*_ reflects the strength of the interaction. *f* (*t*)_*i*_ is a latent variable that represents the normalized proportion of cell type *i* at time *t* (**Fig. 1b**). In constructing the prior for *W*, we penalize self-interactions *W*_*i,i*_, such that *W*_*i,j*_ can be interpreted as the impact of other cell types *j* ≠ *i* on the dynamics of cell type *i*. We justify why this interpretation is reasonable, given our choice of priors below.

#### Prior Distribution over ℱ

Intuitively, we aim for the latent features to align closely with standardized cell type proportions so that *W* (*t*), with suitable restrictions on the diagonal, learns to capture the interactions between cell types by predicting the standardized proportion of one cell type given the standardized proportions of the rest. We achieve this by first defining for every feature an auxiliary zero-mean Gaussian process (GP) on which we perform inference. Formally, for every *k ∈* {1, …, *K* } we define an auxiliary GP with covariance function *K*^*f*^ given by:

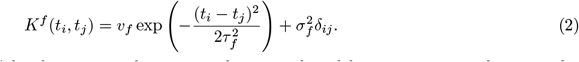

where *δ*_*ij*_ is the Kronecker delta function, and *v*_*f*_, *τ*_*f*_, and *σ*_*f*_ are shared hyperparameters denoting the variance of the latent features, the length scale of the latent features and the noise of the latent features, respectively. We then update this auxiliary GPs using the observations *ŷ*_*k*_ (*t*_*i*_) at the observed time points *t*_*i*_ *∈* 𝒯_*o*_ which provides us with a posterior Gaussian Processes 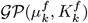that we use as the prior for the Gaussian Process corresponding to feature *f*_*k*_(*t*) in Algorithm 1.

#### Prior Distribution over 𝒲

To further limit the solution space, and improve model robustness and interpretability, we set two constraints on the sampling process of *W*. First, we set off-diagonal elements to zero if the cluster pairs do not express any complementary receptor-ligand pairs and second, we zero out the diagonals^15^. Formally, we achieve this by sampling *W* (*t*) so that for *k, k*^*′*^ *∈ {*1, …, *K}*.

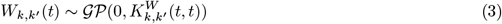

where

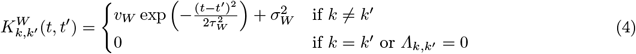

and *v*_*W*_, *τ*_*W*_, and *σ*_*W*_ are shared hyper-parameters, retaining the same interpretations as before. To construct *Λ*, we scan for complementary expression of known RL pairs, i.e. when the sending cluster expresses a ligand at a defined threshold and the receiving cluster expresses the complementary receptor at a similarly defined threshold (**Fig. 1d**). The expression threshold can be set by users and may be datatype-specific. A very high threshold leads to a prior that is too sparse and may lead to losing potentially relevant interactions, whereas a very low threshold may lead to spurious interactions.

We first quantify the number of these interactions expressed by cluster pairs. The number of RL pairs does not necessarily determine the strength of interaction, and a single complementary RL pair could be biologically important. We thus binarize the obtained RL interaction matrix to allow for more flexibility in the model while also limiting the solution space to exclude clusters with no complementary RL expression. This final binary matrix is defined as *Λ*, and as shown in 4, the value of *Λ*_*k,k*_*′* determines whether the model allows for a possible interaction between cell types *k* and *k*^*′*^.

### 2.4 Inference

Due to the complex nature of the model, it is not possible to perform inference analytically. Instead, we deploy a two-step approximate inference method combining both analytic and approximate techniques.

Specifically, we: (1) perform approximate inference to obtain samples from an estimate of the posterior distribution *p*(𝒲_*o*_, ℱ_*o*_ 𝒴_*o*_); and (2) perform ancestral sampling and standard GP conditioning to obtain an approximation to the posterior distribution *p*(𝒲_*u*_, ℱ_*u*_, 𝒴 | 𝒲_*u*_ ℱ_*o*_, 𝒴_*o, o*_). Appendix A provides a justification for this approach. Below we describe in detail how we perform each of these steps.

**Approximate Inference**. To approximate *p*(𝒲_*o*_, ℱ_*o*_| 𝒴_*o*_) we use variational inference implemented using Pyro [4], a probabilistic programming language written in Python. Within this framework, we define a parametrized distribution *q*_*ϕ*_(𝒲_*o*_, ℱ_*o*_) and then optimize the parameters *ϕ* to maximize an estimate of the evidence lower bound (ELBO) [5] via a gradient descent algorithm. In the situation where the hyper-parameters are fixed, this is equivalent to minimizing the KL divergence KL (*q*_*ϕ*_(𝒲_*o*_, ℱ _*o*_) *p*(𝒲_*o*_, ℱ_*o*_| 𝒴_*o*_)). In this case the ELBO is given by

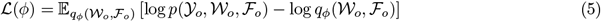

and we define *q*_*ϕ*_(𝒲_*o*_, ℱ_*o*_) as a multivariate Gaussian factorized by:

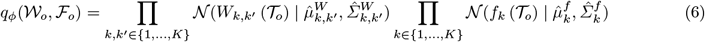

where 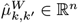and 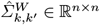are the mean and covariance of 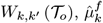and 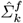 are the mean and covariance of *f*_*k*_ (𝒯), *W*_*k,k*′_ (𝒯_*o*_) denotes the vector of values of *W*_*k,k*_*′* (*t*) for *t∈* 𝒯 _*o*_ and *f*_*k*_ (𝒯_*o*_) denotes the vector of values of *f*_*k*_(*t*) for *t ∈* 𝒯_*o*_.

In other words, we assume that the latent features and the interaction matrix are independent of each other, but they are dependent across timepoints, and this dependency is captured by the mean and covariance of the Gaussian distribution. In practice, we parameterize the Cholesky decomposition of the covariance matrix rather than the covariance matrix itself and use Adam [12] to optimize the parameters *ϕ*. Due to the amount of computation required for using this variational family, we also provide in the package the option to use a mean-field guide where each variable is fully independent from each other and is parameterized by a one-dimensional Normal distribution, where 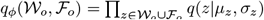.

To perform optimization, we used an expectation with the form 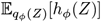where *Z* = (𝒲_*o*_, ℱ _0_) and *h* represents the function inside of the expectation in equation 5. This is problematic for taking the gradient with respect to *ϕ* because it appears both in the distribution with respect to which we are taking the expectation and in *h*. However, using the fact that if *L* = Cholesky(*Σ*), *ϵ* ∼ 𝒩 (0, *I*_*d*_) and *z* = *μ* + *Σϵ* then *z* must be distributed as 𝒩 (*μ, Σ*), it is easy to see that one can rewrite the expectation as 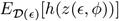where 𝒟 is a multivariate normal distribution with identity covariance and *z*(*ϵ, ϕ*) maps this random variable to the equivalent *Z* values using the above approach. We use this reparametrization trick [13] to construct and estimate the gradient of the ELBO. Appendix B outlines the inference algorithm and provides details of ancestral sampling.

## 3 Experiments and Results

### 3.1 Datasets

#### Simulated Data

We first construct a dataset with known dynamics and interactions to be used for evaluating DIISCO. To define ground truth interactions for W, we simulated 7 non-zero dynamic interactions among 5 cell types using the equations below. To include cases of isolated cell types that do not interact with others, we also allow non-zero diagonals in the prior for a subset of cell types. This can alternatively be captured with a bias term. In particular, we assume:

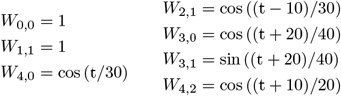

Any interactions not listed above were set to 0. Data points were generated using the following equations:

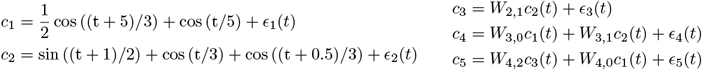

where *ϵ* is a noise term drawn for each time point for each cluster, and *ϵ*_*i*_(*t*) represents noise processes with a standard deviation equal to 0.1. We create the dataset by sampling 100 points uniformly between 0 and 100 and dropping every data point independently with a probability of *p* = 0.2. Based on this generative process, *c*_1_ and *c*_2_ are independent and act on *c*_3_, *c*_4_, and *c*_5_ with different interaction strengths dependent on a time-varying *W*. We set the following elements of the prior matrix:

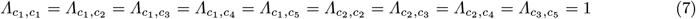

All other elements of *Λ* are set to 0. Importantly, to address potential issues with co-linearity, any cluster pairs that were correlated but not causally related through the defined equations were set to 0. We follow the guide in Appendix D. For fitting the model we let *τ*_*F*_ = 5, *σ*_*f*_ = 0.1, *v*_*f*_ = 3, *σ*_*w*_ = 0.01, *v*_*w*_ = 20, *σ*_*y*_ = 0.1 and we place a gamma hyperprior on the lengthscale of *W* with 90 percent of the mass between 10 to 30. For this simple example, we use the fully factorized guide and use Adam [12] with a learning rate of 0.005

#### CAR-T and MEC1 co-culture

To demonstrate an application of DIISCO on biological data, we collected new single-cell data from an *in vitro* experiment, as a controlled setting. Specifically, we cultured GFP-transduced Chimeric Antigen Receptor (CAR) T cells together with MEC1 cells [24] - a chronic lymphocytic leukemia cell line expressing *CD19* - as the CAR-T cell target (see Appendix C for details). We confirmed dose response activation (cytotoxicity) of T cells by quantifying the percentage of MEC1 cells, showing a reduction ranging from 50 *–* 60% with an effector-to-target ratio of 0.125 to *<* 5% at an effector-to-target ratio of 4, confirming the ability of T cells in targeting and killing the cancer cells. We performed 4 biological replicates with scRNA-seq profiling at 10 time points spanning 24*hrs*, obtaining high-quality data for 49, 283 total cells.

We preprocessed this data by defining major cell types from the combination of all replicate experiments to increase statistical power. Using clustering and expression of curated gene sets, we identified 4 major cell types (metaclusters): cancer cells, exhausted *CD8* ^+^ T cells, activated *CD8* ^+^ T cells, and other *CD8* ^+^ T cells showing neither activated nor exhausted markers. MEC1 cells were annotated based on the expression of *CD19* and *CD79A*. T-cells were annotated based on the expression of *CD3E* and *CD3D*. Activated T cells were defined by expression of *CD69, CD27*, and *CD28*, while exhausted cells were defined by expression of *TIGIT, PDCD1, TGFB1*, and *LAG3*. Clusters that were positive for both T cell and MEC1 cell gene markers were removed as doublets. We acknowledge that doublets could also be containing interacting cells, however, since we only obtain one measurement of receptor or ligand gene, our current method (as well as existing methods) cannot resolve them. Future extensions deploying mixture models could help include doublets.

After cell type annotation, proportions were calculated at each time point, and DIISCO was trained on each experiment separately. To construct *Λ*, we first obtained a set of 8, 234 literature-curated receptor-ligand pairs from OmniPath, a database of cell signaling prior knowledge [26]. For each cell type pair *k, k*^*′*^ in the experiment, we quantified interactions as the number of differentially-expressed ligand genes in cell type *k* with their corresponding differentially-expressed receptor genes in cell type *k*^*′*^ at each time point. These interaction count values were averaged across all time points in the experiment and then thresholded to obtain the binary interaction prior matrix *Λ*, where *Λ*_*k,k*_*′* signifies whether cell types *k* and *k*^*′*^ can interact apriori. The threshold was chosen with a data-driven approach according to the knee-point of sorted values.

### 3.2 Performance on Simulated Data

Evaluating the performance of DIISCO on simulated data, we observe an accurate modeling of temporal dynamics for 5 simulated clusters, even for cell types with complex dynamics and with non-uniform sampling of timepoints (Fig. 3a). Additionally, we find close agreement between inferred interactions and the ground truth specified in section 3.1. Through these simulations, we noted DIISCO’s ability to uncover a wide range of interactions, including constant interactions, monotonically increasing or decreasing interactions, as well as transient interactions (**Fig. 3b**). These various forms of interactions are important to resolve in biological data, especially in studying the impacts of treatments and disease progression, and resolving the timing of critical crosstalk. With treatment administered at discrete time intervals, transient interactions (e.g. *C*5← *C*3 in **Fig. 3b**) may occur that would be missed with static models. In the case of disease progression, interactions between cell types may steadily increase or decrease over time (such as in *C*5 ←*C*1 and *C*3← *C*2), especially as malignant cells become more prominent or more immune cells are recruited to the area. Recapitulating these complex interactions by DIISCO demonstrates its potential to derive insights into dynamic cell communication.

**Fig. 3.**
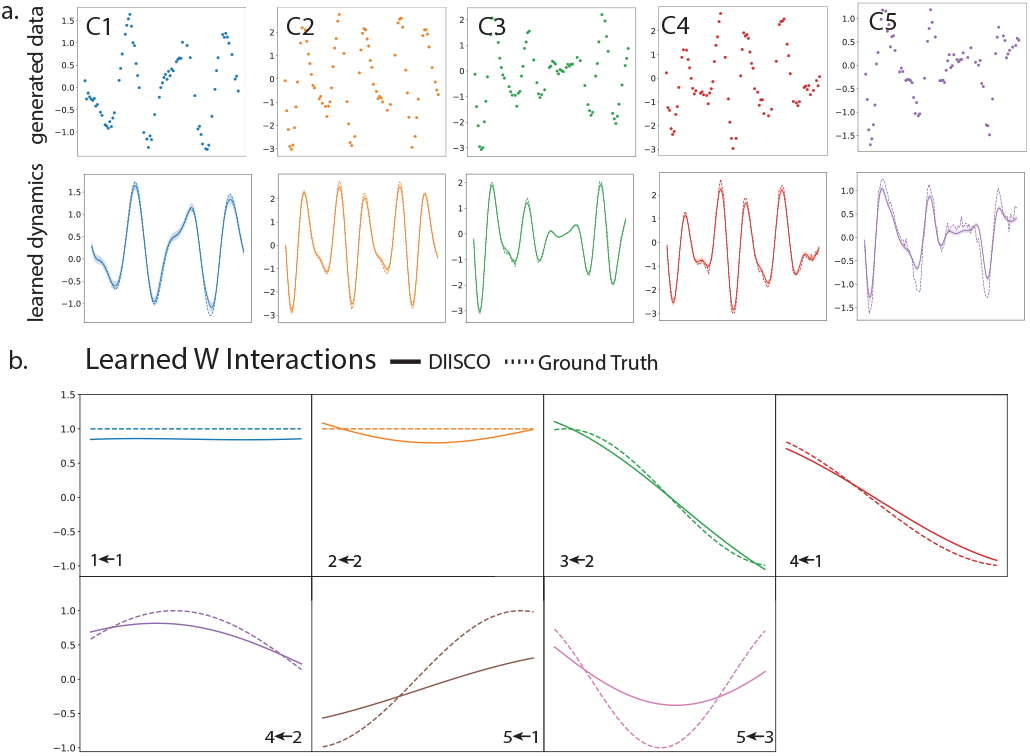
Simulated data and results. **(a)** Simulated data for 5 cell types generated using the process described in section 3.1. Non-uniform sampling of timepoints increases complexity. Top: datapoints for each of the 5 cell types. Bottom: learned DIISCO dynamics after fitting the data. No normalization was used. **(b)** Interactions inferred by DIISCO between the cell types shown in (a). Ground truth W interactions are shown in dotted lines while DIISCO predictions are solid lines. Only dynamic interactions with W (t) > 0.1 in at least one time point t for either true W or learned W are shown.

### 3.3 Analysis of CAR-T and MEC1 Interaction

Applied to CAR-T and MEC1 data, DIISCO learns the proportions of cell types, highlighting the expected decrease in cancer cells over time and the increase in exhausted T cells (**Fig. 4a**). We also infer interactions between cell types and specifically observe strong negative interactions predicted between cancer cells and exhausted CAR-T cells, as well as between activated and exhausted T cells (**Fig. 4b**). Investigating the dynamics of interactions *W* (*t*) over time, as expected, we predict an increase in the strength of the interaction (absolute value of *W*) with co-culture time (**Fig. 4c**) between exhausted T cells and MEC1 cells, as well as in exhausted T cells interacting with activated T cells, which is directionally specific. The former recapitulates the targeting of cancer cells by T cells and the latter may be due to phenotypic shifts from activated cells becoming more exhausted over time [3].

**Fig. 4.**
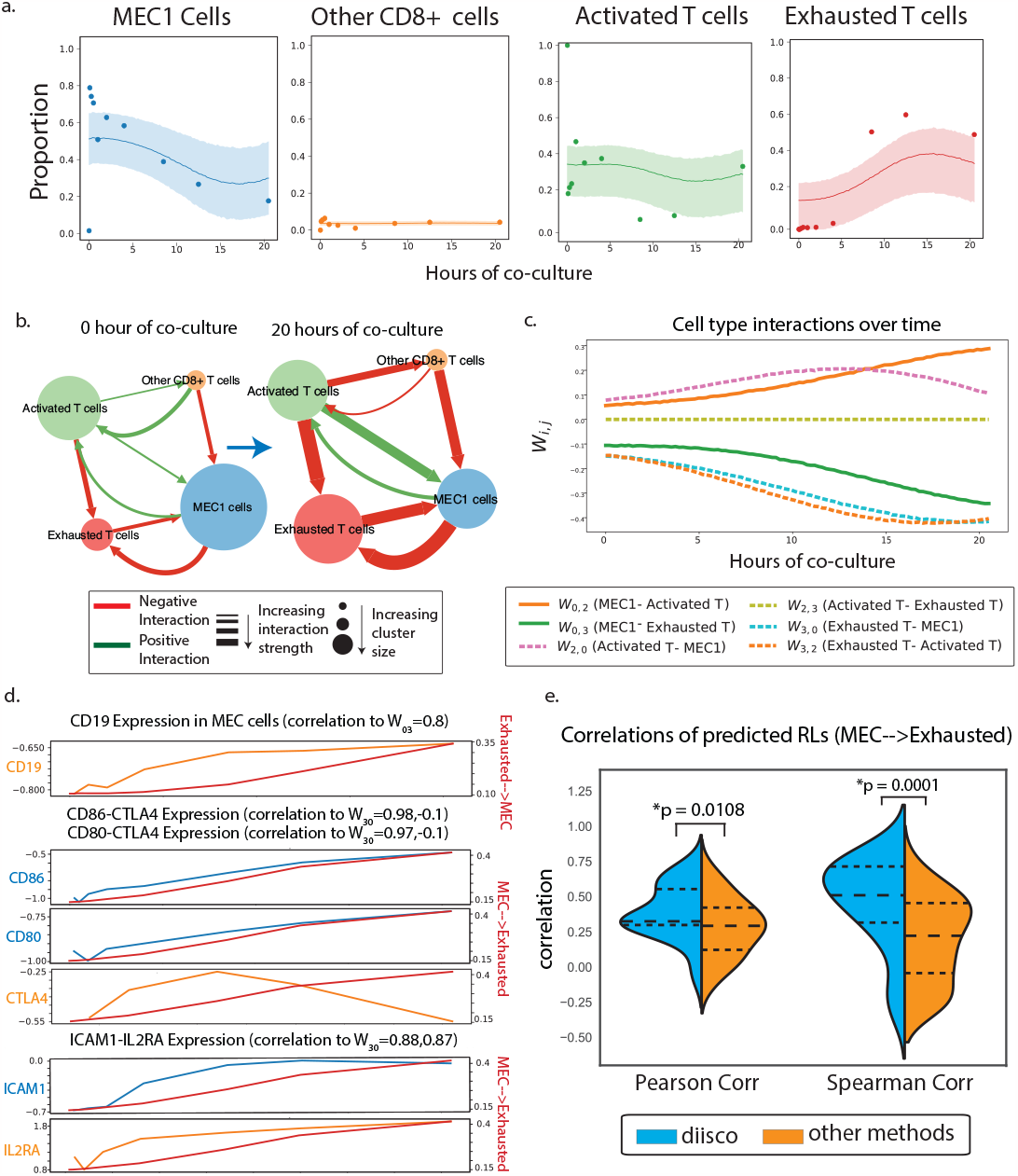
DIISCO performance on CAR-T data. **a**. Temporal dynamics of cell types. Data points represent the measured proportion of cell types over time, and DIISCO output is shown as inferred mean (solid line) and 95% confidence intervals (shaded area) of y for each cell type. **b**-**c** Inferred W interaction matrix at two example time points (b) and over the entire co-culture time window (c). Network node size reflects cell type proportion at that timepoint; arrows and their width designate the direction and strength of inferred interactions respectively. **d**. RL pairs highly correlated with DIISCO predicted W_*Exhausted↔MEC*_ interactions. Expression of ligands is shown in blue, receptors in orange (left axis), and interaction strength in red (right axis). **e**. Comparison of DIISCO predicted RL pairs with those predicted using Cellchat (cc) or CellphoneDB (cdb). Correlations between paired RLs shown for the W_*MEC*→*ExhaustedT cell*_ link. p-values calculated with MWU test.

Importantly, the ability to predict how interactions between exhausted T cells and MEC1 cells evolve with time (**Fig. 4c**) is a unique feature of DIISCO, and other methods are not able to achieve such time-resolved predictions. Indeed, we observe a positive correlation between predicted interaction strength and co-culture time, in line with the cytotoxicity of T cells, i.e. their increased ability to kill cancer cells.

To further investigate mechanisms underlying the interaction between MEC1 cells and Exhausted T cells, we calculated the correlations between different receptor-ligand gene pairs and the learned dynamic interaction (**Fig. 4c**). For calibration, we first confirmed that *CD*19, the target of the engineered CAR protein, is highly correlated with the interaction from Exhausted to MEC1 cells, as expected (**Fig. 4d**). Examining the MEC1→ Exhausted interaction, *CD86-CTLA4* and *CD80-CTLA4* are the top RL pairs ranked by ligand correlation (**Fig. 4d**). *CTLA4* is an established marker of T cell exhaustion, and *CD80* and *CD86* are costimulatory molecules that are expressed in malignant lymphocytes (including B cells) in a number of hematologic diseases [9,27,8]. Ranked by the highest average correlation, *ICAM1-IL2RA* is the top predicted RL pair. ICAM1 is a known regulator of inflammamtory response and is elevated in B-cell CLL patients with more severe disease progression [20,21,7].

Finally, we benchmarked the predicted DIISCO R-L interactions against other methods including CellChat [11] and CellphoneDB [10]. We applied CellPhoneDB and CellChat to log normalized expression data from all time points. We excluded any interactions within the same cluster (i.e. MEC1→ MEC1). RL pairs were filtered based on p-value (*p <* 0.05), and we identified 260 significant pairs using CellPhoneDB and 180 significant interactions using CellChat.

To compare these results to DIISCO-predicted interactions, we calculated the correlation between the predicted receptor and ligand expression over timepoints. Our rationale is that if RL protein complexes mediate cell-cell interaction, their expression levels on the corresponding sender and receiver cell types should change concordantly. If the expression of RL pairs don’t change together over time, then the genes are less likely to be involved in the cellular interaction, and could rather be intrinsic markers of the cell types. We observe a significantly higher Pearson correlation for RLs in DIISCO-predicted interactions than those predicted by other methods (**Fig. 4e**). Since changes in the expression of receptor genes on the receiver cell type may not be linearly associated with the corresponding ligand expression on the sender, we also calculated Spearman correlation, again demonstrating a significant improvement with DIISCO (**Fig. 4e**). Although all 3 methods identify the *CD80/CD86-CTLA4* interactions, only DIISCO predicts the *ICAM1-IL2RA* interaction, which is the interaction with the highest average time varying expression. Our results thus confirm the ability of DIISCO to identify a monotonic increase in the expression of RL pairs underlying cell-cell interactions.

Further examining the overlap between methods, we note that the predicted interactions shared between DIISCO and CellChat are the most correlated with both Spearman correlation and mutual information as another metric, while the interactions predicted by both DIISCO and CellphoneDB are most correlated using Pearson test (**Fig. 5J**). Combined, these results confirm DIISCO’s ability to identify cell-cell communication that can be supported by co-expressed RL genes, from time-series single-cell data.

**Fig. 5.**
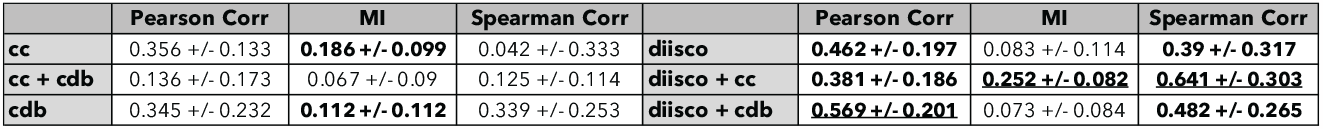
Evaluating temporal correlation of predicted RLs using each method: DIISCO, CellphoneDB (cdb) and Cellchat (cc). A + B refer to RL pairs predicted by method A and B. Table shows 3 different metrics calculated for all R-L pairs: Pearson correlation, mutual information (MI), and Spearman correlation. mean values +/-std are quantified, with top 3 highest scores in **bold** and highest score underlined.

## 4 Discussion and Future Work

As longitudinal single-cell datasets become more prevalent, we believe DIISCO can provide a rigorous and flexible framework for characterizing cell-cell communication and its temporal variability, aiding in the understanding of disease progression, treatments, and complex biological systems. We demonstrate the performance of DIISCO in simulated data as well as cancer-immune cell interaction data. In particular, we show how the integration of time-series data and prior knowledge of protein complexes in a probabilistic model leads to achieving dynamic intercellular interactions. The ability to infer the sign and change in strength of interactions with time is crucial for studying the impact of perturbations such as treatment, and not attainable with previous methods. In addition, DIISCO provides insight into signaling mechanisms mediating dynamic interactions.

Given the model assumptions and limitations, the number of time points greatly affects prediction quality. However, the advantage of the probabilistic framework is that with very few time points or long time intervals, DIISCO will present high uncertainty for estimates. Importantly, as explained in the Appendix D, DIISCO relies heavily on the interaction prior which determines whether the model is identifiable or not. This consideration is important as multi-colinearity might emerge. In particular, in cases where the hyper-parameters are set incorrectly, it is possible that the latent variables lead to seemingly good predictions without this indicating their validity. We thus provide comprehensive guidance for the choice of hyperparameters according to data quality and sampling rate in Appendix D.

Biologically, DIISCO assumes that cellular interactions lead to shifts in the cell type composition of the system. With no changes in cell type proportions or numbers over time, DIISCO is not an appropriate method to use. This may be the case with tissue-resident cells, e.g. those lining the gut, whose interactions do not result in changes in cell type composition but rather result more exclusively in gene expression changes [19].

Similarly, spurious correlations may emerge when using proportions with these types of cell populations due to the constraint of having all points in the simplex. Appendix E includes a short discussion on this issue.

Future application of DIISCO to clinical datasets can elucidate cell-cell communication underlying response or resistance to treatments such as immunotherapy. Deciphering the mechanisms of response and lack of response to therapy allows for more targeted approaches and importantly offers the opportunity to reverse engineer new therapies.

We envision multiple directions for building upon this work. In particular, for clinical samples, we aim to incorporate priors according to sample size (total cells) to improve robustness by down-weighing samples with very few cells. Recent advances in spatial transcriptomics also offer an opportunity to expand DIISCO and infer interactions from both temporal and spatial dynamics by integrating spatial co-localization information. Additionally, DIISCO currently ignores the original space on which the data lies in and makes the simplifying assumption that *y*(*t*) can be treated as a vector in R^*k*^ rather than on the simplex or the space of positive integers. Although these assumptions aid greatly in the interpretability of the latent variables, further improvements using distributions that truly match the real support of the data would lead to better uncertainty estimates and predictions.

## 5 Acknowledgements

We are thankful to David Blei and Dana Pe’er for helpful feedback and discussions. This work was made possible by support from the National Institute of Health (NIH) NCI grants R00CA230195, P01CA229092, Leukemia & Lymohoma Society grant SCOR-22937-22, and the MacMillan Family and the MacMillan Center for the Study of the Non-Coding Cancer Genome at the New York Genome Center. C.P. was supported by the Columbia University Kaganov Fellowship. K.M. was supported by the Richard K. Lubin Family Foundation Fellowship.

## Supplementary Material

### A Justification of Inference Algorithm

According to our model, we are interested in the posterior

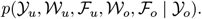

Although, it is not possible to tractably compute or sample from this distribution, we can use its structure to obtain a reasonable approximation. First, using the chain rule of probability, we have:

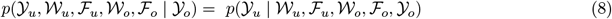

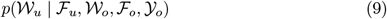

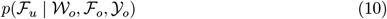

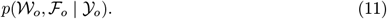

However, by looking at figure (1) we see that in this factorization some dependencies are irrelevant. In particular, we note that the observations 𝒴_*o*_ are independent of everything else given 𝒲_*o*_ and ℱ_*o*_ and therefore, (8) can be written as *p*(𝒴_*u*_|𝒲 _*u*_, ℱ _*u*_), that conditioned on 𝒲_*o*_, ℱ_*u*_ is independent of everything, so (9) can be written as *p*(𝒲_*u*_ 𝒲_*o*_), and that a similar relationship holds between ℱ_*u*_ and ℱ_*o*_, so (10) can be written as *p*(ℱ_*u*_|ℱ _*o*_).s

Using these simplifications, we have:

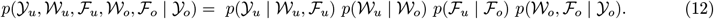

Consequently, if we can obtain a good approximation to the last term, and the first three terms on the right hand side are tractable to compute, we can obtain a good approximation to the full posterior by performing ancestral sampling where we first sample from our approximation *p*(𝒲_*o*_, ℱ_*o*_ | 𝒴_*o*_) and then condition *p*(𝒴_*u*_ | 𝒲_*u*_, ℱ_*u*_), *p*(𝒲_*u*_ | 𝒲_*o*_), and *p*(ℱ_*u*_ | ℱ_*o*_). In the next sections, we describe how we obtain an approximation to to *p*(*W*_*o*_, *F*_*o*_ | *Y*_*o*_) and provide a brief description of how we perform ancestral sampling.

### B Inference Algorithm Details

The detailed inference algorithm is shown in Algorithm 2.4.

**Ancestral Sampling**. To perform ancestral sampling, we execute the following steps:

1. Sample 𝒲_*o*_ and ℱ_*o*_ from *q*_*ϕ*_(𝒲_*o*_, ℱ_*o*_).
2. Compute the posterior distribution *p*(𝒲_*u*_ | 𝒲_*o*_) using the samples from step 1 using algorithm 2.1 from[22] and sample 𝒲^*u*^ from it.
3. Compute the posterior distribution *p*(ℱ_*u*_ | ℱ_*o*_) using the samples from step 1 using algorithm 2.1 from[22] and sample ℱ from it.
4. Compute the posterior distribution *p*(𝒴_*u*_ | 𝒲_*u*_, ℱ_*u*_) 𝒴_*u*_ using equation (1) and sample from it.
5. Return 𝒴_*u*_, 𝒲_*u*_, ℱ_*u*_, 𝒲_*o*_, and ℱ_*o*_.

In practice, since steps 2 and 3 are computationally expensive due to the computation of the posterior of a Gaussian Process, we sample *p*(𝒲_*u*_ | 𝒲 _*o*_) and *p*(ℱ_*u*_ | ℱ _*o*_) multiple times per sampling of _*o*_ and ℱ_*o*_ respectively. The inference algorithm is detailed below.

### Additional Practical Considerations

During training, we use early stopping by defining an epoch as 1000 iterations of the optimization algorithm and stopping when the ELBO has not increased for 10 epochs. For hyper-parameter selection, we follow the recommendations detailed in Appendix D in the appendix but set a hyper-prior on the length scale of *W* (*t*) to allow for flexibility in the model. To infer this value we augment the variational family above with an additional term 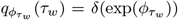 where *δ* is the delta distribution. As further discussed in Appendix D we emphasize that choosing these hyper-parameters is crucial for the model to adequately perform its function as incorrectly setting these values can lead to degenerate solutions with non-identifiability.

#### Algorithm 2

Simplified Inference Algorithm used by DIISCO

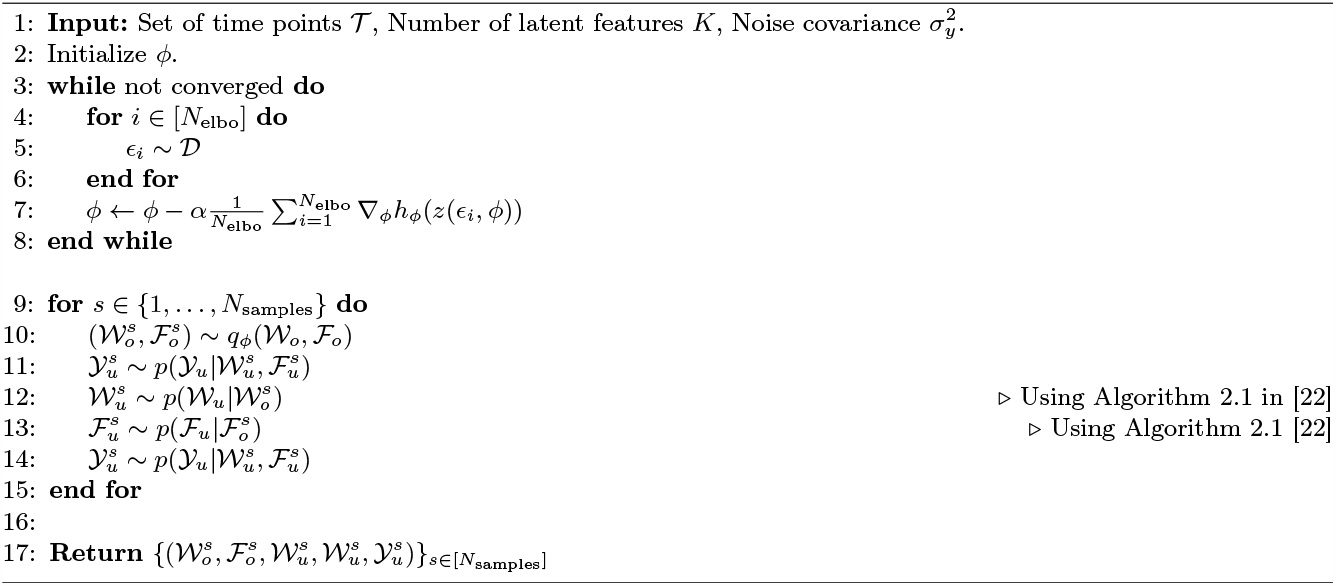

### C Details of CAR-T Experiment

CD19 CAR-T cells were generated by transducing healthy donor T cells with 3rd generation lentiviral vectors encoding a bicistronic construct containing either FMC63 CD19 scFv-CD28-CD3z and green fluorescent protein (GFP) or FMC63 CD19 scFv-41BB-CD3z. Peripheral blood mononuclear cells were re-suspended at 2 *×* 106/ml and seeded at 1ml per 24 well plate and activated with CD3/CD28 Beads. The next day fresh media was added with IL-2 to a final concentration of 100*IU/ml* and 6 hours later cells were harvested, counted and re-suspended at 0.6 *×* 10^6^*/ml* and 0.5*ml* was seeded into a 24 well plate pre-coated with retronectin. 1.5*ml* of lentiviral supernatant was added to each well with fresh IL-2 to a final concentration of 100IU/ml and spun for 40 minutes at 1000G. Two days later cells were harvested, re-suspended at 0.5*×* 10^6^*/ml* with 50*IU/ml* of IL-2 and left to expand and split every 48 hours. Transduction efficiency was assessed by determining the percentage of GFP+ T cells.

For co-culture experiments co-cultures of CD19 CAR-T cells and MEC1 cells, a CLL cell line that constitutively expressed CD19 were established at various effector-to-target ratios and at different time points (as detailed in the method). Co-cultures were harvested together and stained with hashing antibodies (Biolegend), normalized and prepared for single-cell RNA sequencing on the Chromium 10X platform.

### D Hyper-parameter Selection Guide

Choosing the adequate hyper-parameters is crucial for the success of the model. In particular a bad selection of hyper-parameters can lead to non-identifiability or results that don’t align adequately with the interpretation.

In this section we provide a summary of the most relevant hyper-parameters of the model, their interpretation, and suggestions and reasoning for how to set them.

Table 1 contains a summary of the hyper-parameters used by the model. We will describe below the role that each of these hyper-parameters play and how to set them. We will go from most important one to least important one:

**Table 1.** Hyperparameters and their Descriptions.

| Symbol | Description |
| --- | --- |
| $\tau_f$ | Lengthscale for $f$, controls how flexible is the prior over the latent features. |
| $\tau_w$ | Lengthscale for $W$, controls how flexible is the matrix and how much information is shared across time points. |
| $v_f$ | Variance for $f$, controls the magnitude of the latent features. |
| $v_w$ | Variance for $W$, controls the magnitude of $W$ matrix |
| $\sigma_f$ | Standard deviation for $f$, controls the amount of non informative noise we believe is in the latent features. |
| $\sigma_w$ | Standard deviation for $W$, controls the amount of non informative noise we believe is in $W$ and serves mostly as a stability parameter during optimization. |
| $\sigma_y$ | Standard deviation for $y$, controls the amount of noise the model assumes is in the data. |

1. Lengthscale *τ*_*w*_: This is by far the most important parameter of the model. It controls the flexibility and smoothness of the *W* matrix which determines both how much information is used to inform the value at a predicted time point and how quickly this value changes. Ideally, one would like to set it from domain knowledge but it can be set with intuition derived from the data as follows. Intuitively, two points a distance of a length-scale away have correlation of ≈ 0.6 ≈1*/*2 where this correlation is measured with respect to random function draws. A good rule of thumb is that the length scale should be roughly at least as big as the maximum distance in the data between the largest and the smallest of any *j* sequential data points, where *j* is the largest number of non-zero entries in a row of *Λ*. Mathematically,

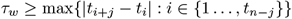

where

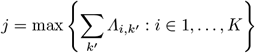

Consequently, the sampling frequency of the data should be such that this length-scale is adequate to model the flexibility we expect in the *W* matrix. This is not a rule applicable everywhere but it adheres to the intuition that our model is performing a form of approximate local linear regression and this is the minimum number of points we would need for such a scheme to work approximately, taking into account the fact that a lot of information is being shared.
2. Lengthscale *τ*_*f*_ : This plays the normal role of the length-scale in traditional Gaussian Processes and can be set so that the prior matches the intuition of the user about the latent functions, or using one of the traditional methods implemented in any package for setting this hyper-parameter.
3. Variances *v*_*f*_, *v*_*w*_: These values play the same role as the variance in a standard Bayesian linear regression and can determine the magnitude of the functions drawn. In our case when standardizing we set them to values slightly above one for *f* and higher for *v*_*w*_ to indicate a weak prior. As *v*_*f*_ affects the covariance, it should be handled jointly with *τ*_*w*_ to express beliefs about the flexibility of the functions.
4. Noise *σ*_*f*_ and *σ*_*y*_: Indicate how much noise we believe is in our observations. The higher the noise, the more points the model will need to change the latent distribution. In our case, we used values lower than one to indicate low noise in our observations.
5. Noise *σ*_*w*_: In our case, this is mostly an optimization stability parameter that can be set very small and is only useful to avoid numerical problems. We set it to 0.001 in our experiments and can be left to this default value.

### E Assumptions for Application to Cell Type Proportions

Our model is designed to work equally with proportions as well as raw count data. However, one must have particular care when working with proportions to make sure that the assumptions of the model are met.

As detailed by Aitchison [1], compositional data, i.e data points that lie on the simplex, present a particular challenge when thinking about their correlation structure and what it implies about the real biological process.

In particular, if we assume that our proportions *y*(*t*) *∈* Δ^*K*−1^ emerge from some real process with some absolute number of counts 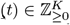 and it is the case that Σ_*k*_ *c*_*k*_(*t*), i.e the total number of counts, varies widely through time then it is possible that the correlations that the model learns will not be meaningful. This is a limitation not only for this model but also for any model that only uses proportions to understand the relationship between the variables. If however, it is the case that Σ_*k*_ *c*_*k*_(*t*) ≈*C* for all *t* the inferences made by the model will be valid.

We demonstrate this issue with the following example. Assume that we have two clusters with the following dynamics:

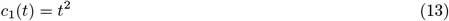

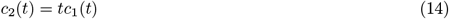

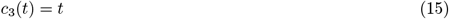

And with proportions

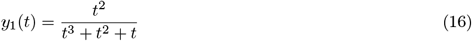

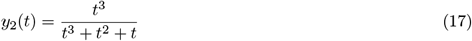

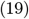

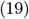

Clearly Σ_*k*_ *c*_*k*_(*t*) = *t*^3^ + *t*^2^ + *t* is not constant. If one were working with raw values, we would like to say that *c*_1_, *c*_2_ positively interact in that they increase simultaneously. However, when considering proportions, the interpretation is different because now as *y*_2_ is increasing, *y*_1_ is decreasing which would suggest a negative interaction.

We thus advise using DIISCO in settings where the total number of cells across time points does not have extreme variability or to drop low-quality samples.

## Footnotes

13 We also allow for a bias term b(t) such that *Y (t) = W* (t)f*(t) + b(t) + ϵ(t)* where b(t) is centered at 0 and has a kernel like that of W (t) but without any interaction regularization. For brevity, we omit it as it is equivalent to extending the model dimension by one, ignoring the last coordinate of y(t), and setting the last coordinate of f(t) to be a GP centered at 1 with infinite length-scale.

14 The arrows 𝒲_o_→ 𝒲_*u*_ and ℱ_*o*_ → ℱ_*u*_ represent tractable distributions that can be computed analytically and can be sampled due to the properties of Gaussian processes. Thus, the model only requires tractable operations that lead to the joint distribution described.

15 In the case when we are dealing with proportions and not raw counts this also ensures that we avoid a trivial solution due to theΣ_*k*_ y_*k*_(t) = 1.

